# Efficient searches in protein sequence space through AI-driven iterative learning

**DOI:** 10.1101/2024.12.31.630868

**Authors:** Ignacio Suarez-Martin, Valeria A. Risso, Rocío Romero-Zaliz, Jose M. Sanchez-Ruiz

## Abstract

Protein sequence space is vast. This fact, together with the prevalence of epistasis, hampers the engineering of novel enzymes through library screening and is a major obstacle to any attempt to predict natural protein evolution. Recently, specialized methodologies have been used to determine fitness data on ∼260000 sequences for the gene of the enzyme dihydrofolate reductase and antibody affinity data for all combinations of the mutations present in the receptor binding domain (RBD) of the Omicron strain of SARS-CoV-2 (∼30000 variants). We show that, upon iterative training on a total of just a few hundred variants, various state-of-the-art AI tools (multi-layer perceptron, random forest and XGBoost algorithms) find very high fitness variants of the enzyme and predict the antibody-evasion patterns of the RBD. This work provides a basis for efficient, widely applicable, low-throughput experimental approaches to assess viral protein evolution and to engineer enzymes for biotechnological applications.

---

The protein sequence space is the mathematical space of all possible sequences. The full space for a given sequence length (L) is enormous: 20^L^ amino acid sequences encoded by 4^3L^ gene sequences. Searching the sequence space for useful or relevant protein properties is a daunting task, in particular because the prevalence of epistasis (*i*.*e*., strong non-additivity of mutational effects) makes it impossible to describe the values of the relevant property for all sequences in terms of a comparatively small number of individual mutational effects.^1-5^ Indeed, laboratory directed evolution, the most generally applicable approach to protein engineering,^6^ is typically sluggish and may require many rounds of effort-intensive library screening to achieve the desired levels of the targeted properties.^7^ This obviously reflects the vastness of protein sequence space and the fact most sequences do not encode for proteins with biotechnologically useful properties. As a result, there has been considerable interest in the development of computational and experimental methodologies to focus directed evolution to small regions of the sequence space that are predicted to be particularly relevant for the targeted protein property.^8-17^ However, while methodologies of guided directed evolution represent an enormous advance for protein biotechnology, they do not fully solve the search problem. Even if some rational approach pinpoints, say, 10-20 mutations that may potentially modulate the targeted protein property, the corresponding combinatorial library may span hundreds of thousands of variants. Specialized methodologies to exhaustively screen such libraries are not generally available, in particular for many of the biotechnologically relevant goals, such as the enhancement of a low-level promiscuous activity or the engineering of enzyme regio- and stereo-selectivity.

Here, we show that efficient searches in regions of the protein sequence space spanning, at least, a few hundred thousand variants can be performed using simple AI-tools that are iteratively trained on a total of only a few hundred variants. The methodology we propose is akin to focused directed evolution, but it does not rely on previously known or predicted information. Rather, it is the AI tool which, upon iterative learning, will focus itself to very small regions of the sequence space that encode for protein variants with the targeted properties. To test our proposal, we have taken advantage of a recent study in which a specialized methodology has been used to determine fitness data on ∼260000 sequences for the gene of the enzyme dihydrofolate reductase.^18^ We will use this data set to address the common biotechnological scenario in which enzyme optimization is essential and a few variants (or even a single variant) with the desired properties suffice to enable a practical application.

The state-of-the-art AI tools we have used (see Fig. 1) are: 1) multi-layer perceptron,^19^ a feedforward neural network (Fig. 1a); 2) random-forest algorithm,^20^ which combines several decision trees to generate a result (Fig. 1c); 3) XGBoost algorithm,^21^ which provides a regularized gradient boost to the random forest (Fig. 1d). When used as regressors, we find that the three AI tools identify very rare variants of highly enhanced fitness, even only after a few rounds of iterative learning involving a total of just a few hundred of sequences, *i*.*e*., a number much smaller than the size of the explored region of the sequence space. This result is particularly remarkable, since analyses of the original data^18^ supported the prevalence of epistasis, including higher-order epistasis (*i*.*e*., non-additivity of the effects of more than two mutations), which brings about a rugged, difficult to search fitness landscape.

**Fig. 1.**
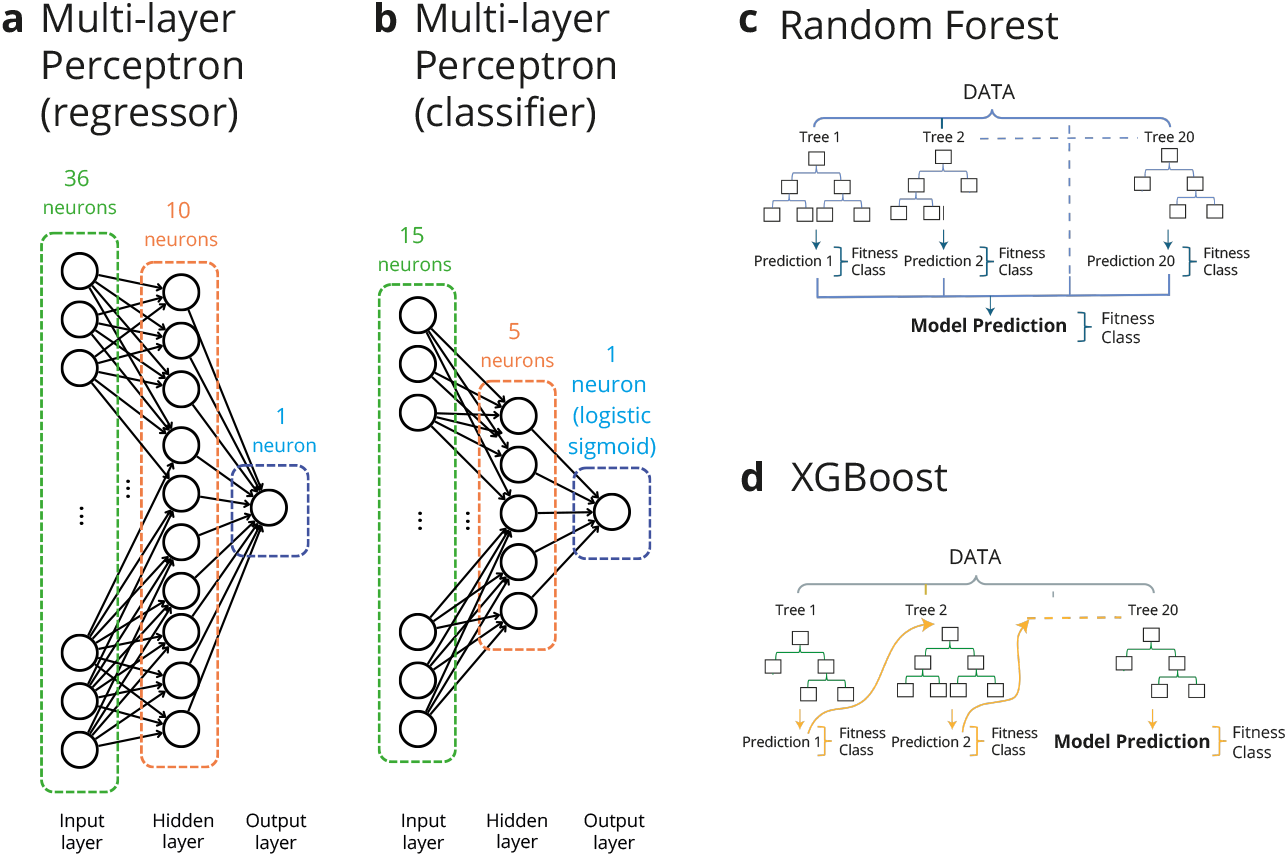
Types of AI tools used in this work. **a**, Multi-layer perceptron used as a regressor to search the sequence space of the enzyme dihydrofolate reductase. Since we use one-hot encoding, there are 36 neurons in the input layer corresponding to the 4 possible nucleotides at the 9 targeted positions in the enzyme gene (see text for details). There is a single, fully connected intermediate hidden layer of 10 neurons. **b**, Multi-layer perceptron used to search the sequence space of the receptor binding domain (RBD) of the SARS-CoV-2 virus. It includes as output a single neuron with a logistic sigmoid activation function, in such a way that it acts as a classifier that allocates variants to two classes: “evasion” and “no evasion”. The input layer has 15 neurons, corresponding to the 15 mutations that separate the RBD of the Omicron BA.1 strain from that of the parent Wuhan Hu-1 strain. There is a single, fully connected intermediate hidden layer of 5 neurons (see text for details). **c**, Random Forest algorithm, which combines 20 decision trees to generate an output. **d**, XGBoost algorithm, which provides a regularized gradient boost to the Random Forest. Note that both, the Random Forest and XGBoost algorithms were used as regressors (to search the enzyme sequence space) and classifiers (to search the RBD sequence space).

Beyond the obvious implications for protein engineering, the fact that the protein sequence space is vast is also a major obstacle to any attempts to predict viral protein evolution on the basis of laboratory studies on the relevant biomolecular interactions. This is so because viruses have a capability to search protein sequence space that currently available experimental approaches cannot match. This capability results, not only from the high viral mutational rates, but also from the huge numbers of virions during an infection. For instance, the number of virions in each person infected by SARS-CoV-2 has been estimated to be 10^9^-10^11^ at peak infection.^22^ Overall, viruses may efficiently search protein sequence space and, when faced with new challenges, they may come up with solutions involving large numbers of mutations. For instance, the original Omicron strain of SARS-CoV-2 evades many antibodies against earlier variants of the virus and has 15 mutations in the receptor binding domain of the spike protein, which is the main target of neutralizing antibodies,^23^ and variants with many additional mutations are emerging.^24,25^ Recently, Moulana et al.^23^ used a specialized methodology to determine antibody affinity data for all combinations of the mutations present in the receptor binding domain (RBD) of the Omicron strain of SARS-CoV-2, *i*.*e*. about ∼30000 variants. In order to test the generality of our approach, we used the iterative learning approach using state-of-the-art AI tools depicted in Fig. 1 to search the RBD sequence space. Here, our analyses aimed at achieving a reliable classification of the variants in terms of antibody evasion versus non-evasion. That is, the AI tools will be used as classifiers (Fig. 1b, 1c and 1d). We find that upon iterative training on a total of just a few hundred of variants, the AI tools correctly assign most variants to the evasion or non-evasion classes for three different neutralizing antibodies. Again, this is a remarkable result, since epistasis plays an important role in determining antibody affinity in this case^23^ and this should bring about a rugged, difficult to search landscape.

## RESULTS

### Searching the sequence space of the enzyme dihydrofolate reductase

Papkou et al.^18^ have recently used CRISP-Cas9 deep mutagenesis to randomize 9 positions in the gene of the enzyme dihydrofolate reductase. The randomized positions correspond to the codons for three consecutive amino acid residues located at the enzyme active site (see Fig. 2a). Randomizing the 9 positions resulted in a library of ∼260000 nucleotide sequences (DNA genotypes). Expression of the library in *E. coli*, exposure of the *E. coli* population to the antibiotic trimethoprim (potentially degraded by active variants of the enzyme), followed by deep scanning mutagenesis, allowed fitness values to be assigned to essentially all variants in the library. Fitness was defined by variant frequency (after selection) in a logarithmic scale. The wild-type enzyme was taken as reference and assigned a fitness value of zero. Fitness values thus calculated reflect contributions from several factors, including the effects of mutations at the three encoded amino acid positions on enzyme activity and proper folding, as well as the effects of the nucleotide sequence on expression levels. Regardless of complex molecular interpretations of fitness in this case, the availability of enzyme fitness values for a library of ∼260000 variants allowed us to test the approach proposed in this work in the context of enzyme optimization.

**Fig. 2.**
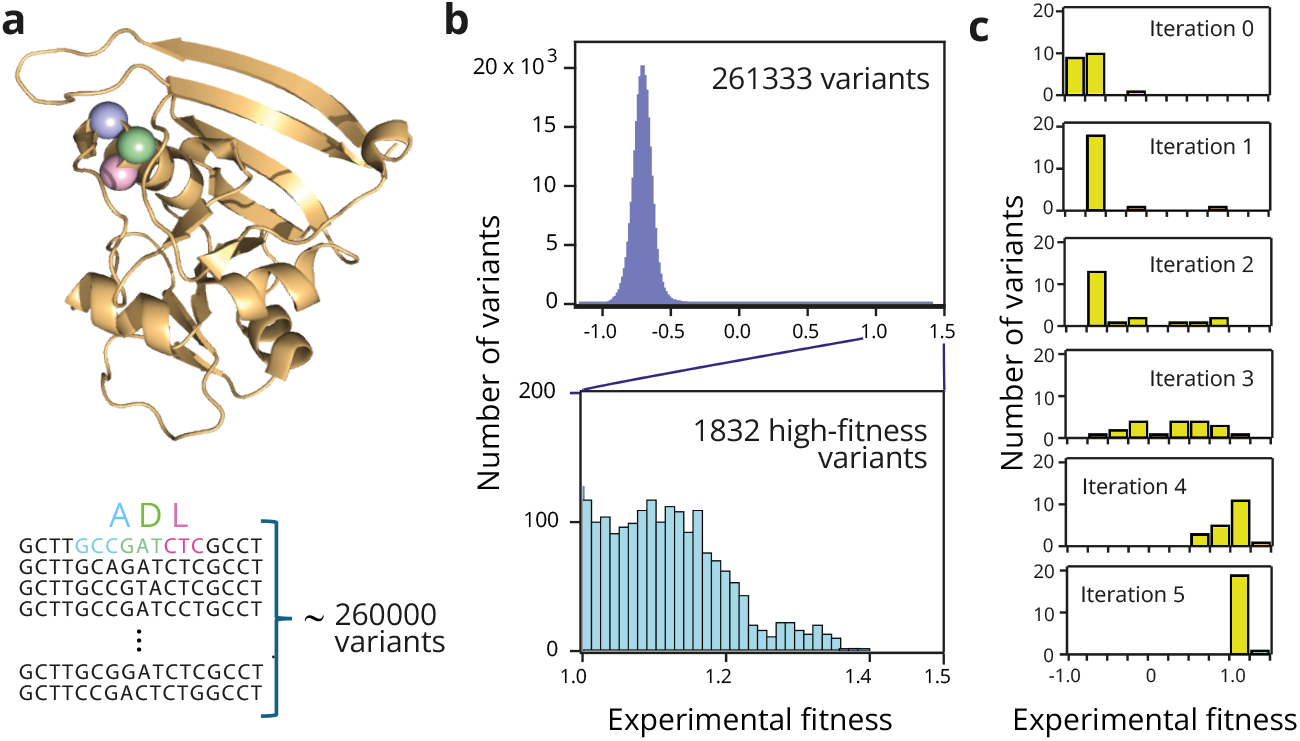
Fitness distribution for variants of dihydrofolate reductase. **a**, 3D-structure of dihydrofolate reductase (PDB ID 6XG5) and section of the enzyme gene sequence including the 9 positions that were randomized^18^ to generate a library of ∼260000 variants. These 9 positions in the gene encode for 3 amino acid residues in the enzyme active site (A, D an L in the wild-type enzyme). The location of the three residues is shown in the 3D-structure. **b**, Distribution of fitness values for the ∼260000 variants generated as shown in a. The logarithmic fitness scale is defined in such a way that the fitness of the wild-type enzyme is zero. Most variants display impaired fitness, while only about 0.7% of the variants have a much-enhanced fitness value above unity (blow-up shown in the lower panel). **c**, Illustrative example of an AI-driven iterative learning search of the library. A multilayer perceptron (Fig. 1a) is initially trained with 20 randomly chosen variants (iteration 0). The sequence/fitness data for the top 20 predictions are then incorporated to the training set for the next round of training, thus initiating the iterative learning procedure. The plots show the distribution of the experimental fitness values for the top 20 variants of the training set at the successive iterations.

Fig. 2b shows the distribution of fitness values for the ∼260000 variants of the combinatorial library. The distribution peaks at negative values of about -0.7 indicating that most variants in the library are substantially worse than wild type (which was assigned fitness value of zero). This result likely reflects that, for many variants, proper folding is compromised, enzyme activity is impaired, or expression levels are greatly diminished. A small fraction of the variants displays enhanced fitness with respect to wild type and only about 0.7 % of the variants have fitness values of unity or above, as shown in the lower panel of Fig. 2b. We have selected fitness>1 as the optimization target for the analyses described below.

The sequence and fitness value of the wild-type enzyme is, of course, known. However, our analyses are meant to represent the common experimental scenario in which no information about the properties of the library variants is *a priori* available. Therefore, we ignored any previous knowledge and started searches by randomly selecting 20 variants. The chosen AI tool (see Fig. 1a, 1c and 1d) was then trained on the sequence/fitness data for these 20 variants and the trained tool was then used to predict fitness values for all variants in the combinatorial space (4^9^=262144 sequences). The sequence/fitness data for the top 20 predictions for which experimental fitness values are available, are then incporated to the training set for the next round, thus initiating the iterative learning procedure. An illustrative example of this protocol is shown in Fig. 2c. Initially, the fitness values are low, reflecting the distribution of fitness values for the whole library (Fig. 2b). Yet, after only 5 rounds of iterative training involving a total of only 120 variants, the 20 top predictions already display examples of sequences that meet our optimization target, *i*.*e*., sequences with experimental fitness above unity.

To explore the extent to which the success of the iterative search process depends on the starting set of 20 variants, we performed, for each of the three types of AI tools used, 1000 replicas of the protocol described above. We focused the analysis of the results obtained on the best variant from each search, in congruence with the common protein biotechnology scenario in which a practical application may be enabled by a single variant with the required enhanced properties. Fig. 3a shows the distribution of the best variants at the fifth round. A substantial fraction of the searches meets the proposed target (fitness higher than unity) at the fifth iteration (*i*.*e*., after training with a total of 120 variants) with the performance of the different types of AI tools following the ranking GXBoost > Random Forest > Multi-layer Perceptron (Fig. 3a). The success statistics is substantially improved if, instead of a single AI tool, we consider ensembles of 5 AI tools of the same type, each one being independently trained. As shown in Fig. 3b, most of the ensembles meet the optimization target (fitness higher than unity) at the fifth iteration, which involves in this case training with a total of 5×120=600 variants. Actually, at the fifth iteration, most of the ensembles achieve fitness values higher than 1.2, which corresponds to a tiny fraction of the total number of sequences considered (Fig. 2b).

**Fig. 3.**
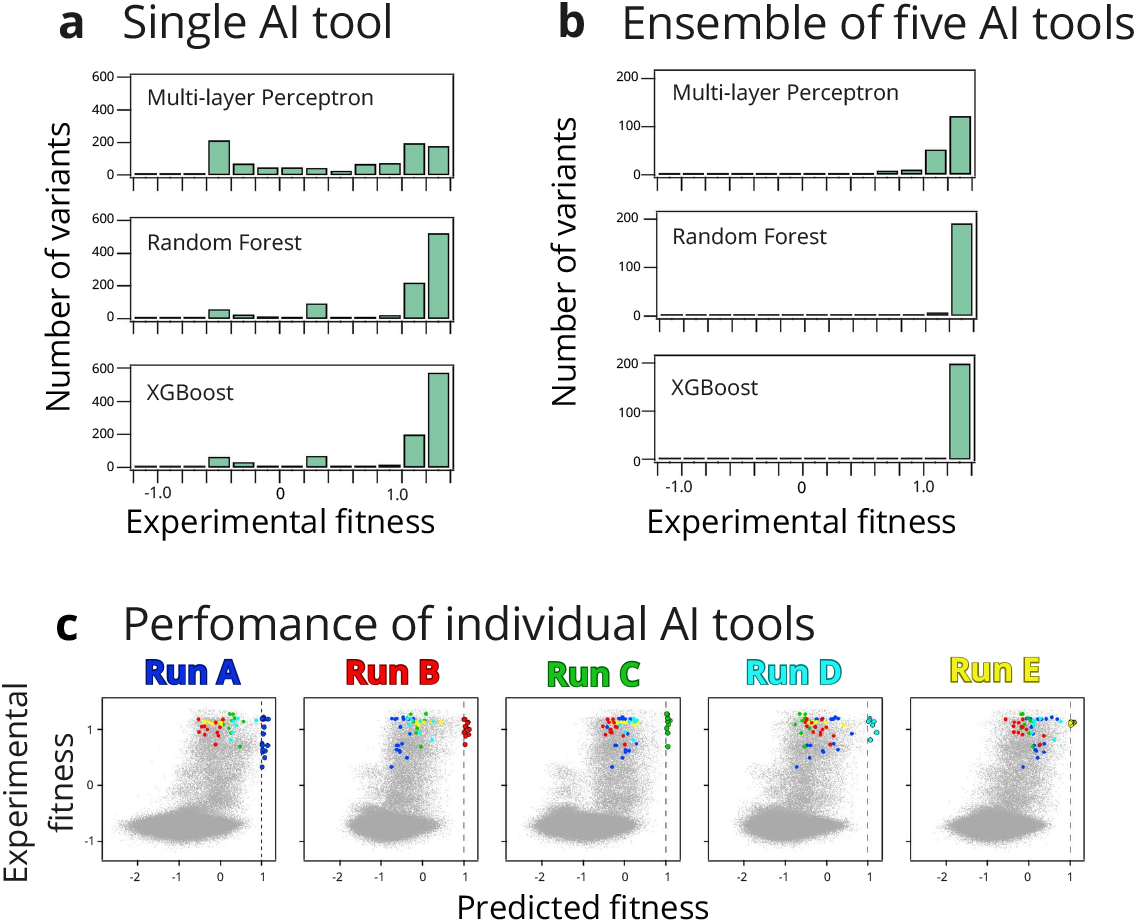
Finding high-fitness variants of dihydrofolate reductase through AI-driven iterative learning. **a**, Distribution of best (higher experimental fitness value) variants at the 5^th^ iteration from 1000 search replicas using three different AI tool (Figs. 1a, 1c and 1d). **b**, Distribution of best (higher experimental fitness value) variants at the 5^th^ iteration for ensembles of 5 AI tools of the same type in each case. 200 replicas of the ensemble search were performed for each AI tool. **c**, individual AI-tools do not achieve an accurate description of the fitness value for the whole library but, rather, they locate different high-fitness regions in sequence space. This is illustrated by the plots of experimental versus predicted fitness for 5 searches based in multi-layer perceptrons at the 5^th^ iterative round. Each run predicts a certain number of high-fitness variants which differ from the high-fitness variants predicted by the other runs. To make this fact visually clear, the high-fitness predictions from each perceptron are colour coded.

Finally, it must be noted that each trained AI tool does not (and it is not meant to) predict experimental fitness for the whole combinatorial library to any significant degree of accuracy. Rather, each AI tool approaches a given peak in the fitness landscape, with different successful tools approaching different peaks. That is, different searches lead to different sets of high-fitness variants, as it is illustrated in Fig. 3c.

### Searching the sequence space of the receptor binding domain of SARS-CoV-2

The receptor binding domain (RBD) of the spike protein provides the major antigenic target of the SARS-CoV-2 virus. 15 mutations separate the RBD of the Omicron BA.1 strain from that of the parent Wuhan Hu-1 (see Fig. 4a). Moulana et al.^23^ generated a complete combinatorial library including all the RBD intermediates between the parent and the Omicron strain, *i*.*e*., 2^15^=32768 amino acid sequences. They displayed the library on the surface of yeast and used flow cytometry coupled to sequencing to determine the binding affinities of several monoclonal antibodies to most of the RBD variants in the library. These included three neutralizing antibodies (LY-CoV016, LY-CoV555, REGN10987), which have been used in the treatment of COVID-19, although the three of them are evaded by the Omicron strain. The availability of antibody dissociation constants for a library of about ∼30000 RBD variants allowed us to test our approach in the context of antibody evasion.

**Fig. 4.**
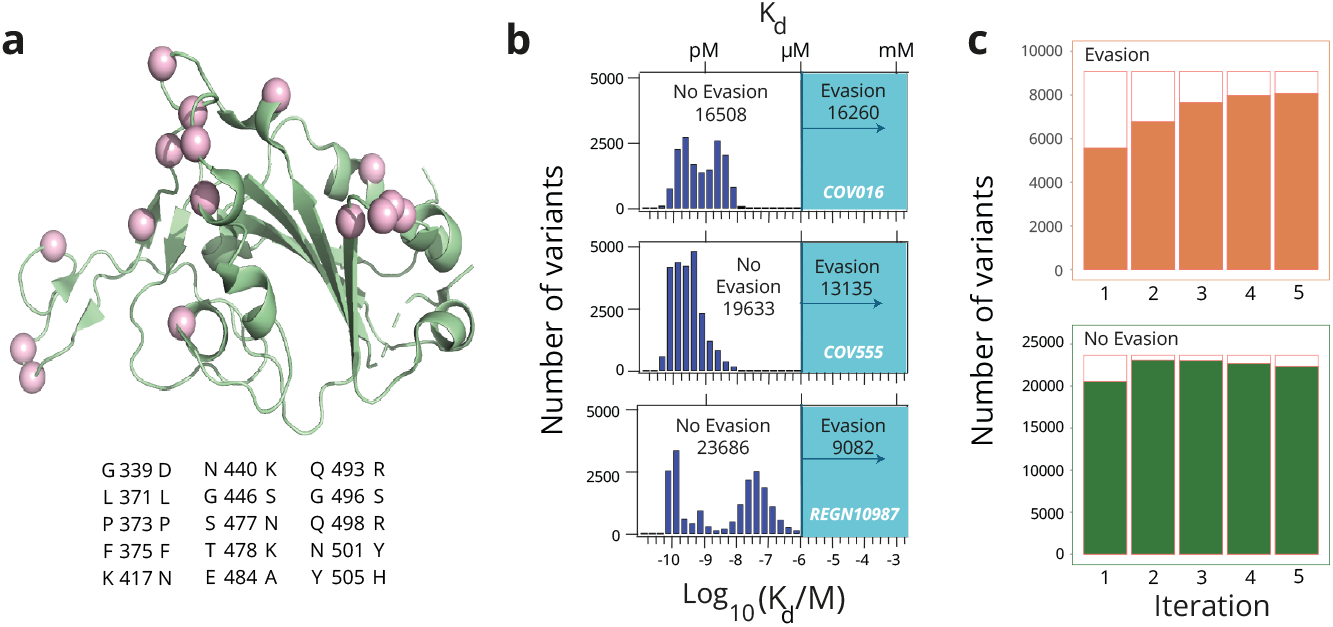
Distribution of antibody affinity to variants of the receptor binding domain of the SARS-CoV-2 virus. **a**, 3D-structure of the RBD (PDB ID 7BH9) showing the positions at which there are amino acid differences between the RBD of the Omicron BA.1 strain and that of the parent Wuhan Hu-1 strain. The specific amino acid replacements are listed below the structure. The combination of these replacements gives rise to a library of 2^15^=32768 variants. **b**, Distribution of antibody-RBD dissociation constants for the variants of the library and three different neutralizing antibodies: LY-CoV016, LY-CoV555, REGN10987.^23^ Dissociation constants higher than 1 micromolar could not be determined experimentally and are taken to define antibody evasion. Numbers of evading and non-evading variants for each antibody are shown. **c**, Illustrative example of an AI-driven iterative learning search of the library. XGBoost (Fig. 1d) was initially trained with 20 randomly chosen variants and predictions were used to increase the training set with 20 additional variants at each iteration round. The plots show the total number evading and non-evading variants for the antibody REGN10987, as well the number of correct predictions in each case (solid colour).

Distributions for the logarithms of the experimental antibody-RBD dissociation constants are shown in Fig. 3b. Many dissociation-constant values are much smaller than 1 micromolar, indicating tight antibody-RBD binding. Yet, for a substantial number of variants, dissociation constants are larger than 1 micromolar and could not be determined with the methodology used.^23^ These variants do not bind the antibodies or, at least, they bind them weakly and likely non-specifically.^23^ We take the range of dissociation constants of 1 μM and above to correspond to antibody evasion (Fig. 3b). Under this definition of evasion, 50 %, 40 % and 28 % of the RBD variants evade the antibodies LY-CoV016, LY-CoV555, REGN10987, respectively (Fig. 3b).

We aimed to show that an iterative learning approach capable of efficiently classifying the sequences in two classes, evading and no evading, for each given antibody. The process was started by randomly selecting 20 variants. The selected AI tool was then trained on the sequence/evasion data for these 20 variants and the trained tool was used to classify all variants in the combinatorial space (2^15^=32768) into the two specified classes. To avoid losing variant diversity in the training set, the 20 variants incorporated in each subsequent round of training were selected to have a minimum number of sequence differences (typically 3: see methods for details) with those in the previous training set. As an illustration of the approach, Fig. 4c shows the result of the training of a XGBoost tool to classify the REGN10987 evasion data. After only five rounds of iterative training (involving a total of 120 variants in the sequence set) most of the ∼30000 sequences are correctly classified into the evasion versus no-evasion classes.

To explore the extent to which the success of the iterative search process depends on the starting set of 20 variants, we performed 500 replicas of the protocol described above for each of the three types of AI tools used (Fig. 1) and with the three antibodies studied (Fig. 4b). The results are displayed as box-and-whiskers plots in Fig. 5a. Even after 5 iterations (a total of 120 variants in the training set) the average success in variant classification (evasion versus no evasion) is higher than 80% and actually approaches 100% in some cases. The performance of the different types of AI tools appears again to follow the ranking GXBoost > Random Forest > Multi-layer Perceptron. Furthermore, it emerges that the efficiency of the search also depends on the antibody considered, and the ranking LY-CoV016 > REGN10987 > LY-CoV555 is visually apparent in Fig. 5a.

**Fig. 5.**
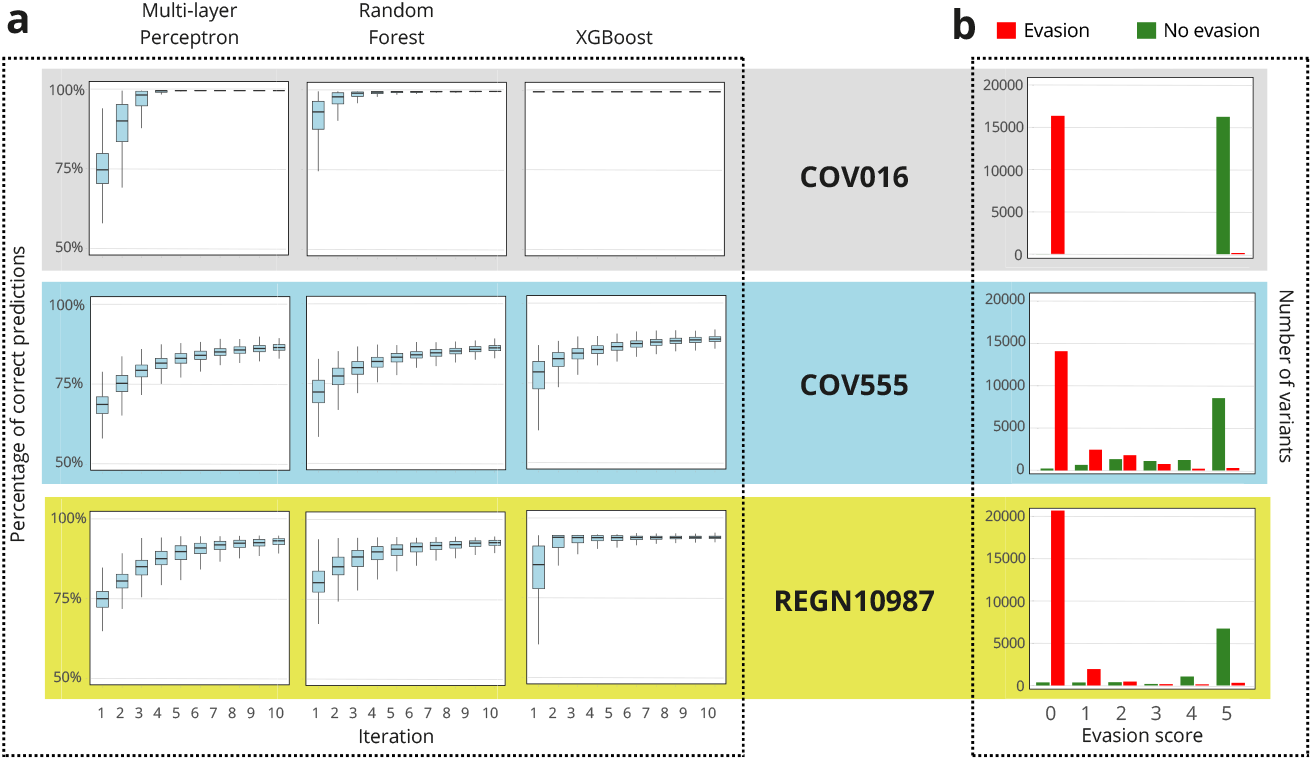
Using AI-driven iterative learning to classify RBD variants into evading and non-evading classes. **a**, For each antibody/AI-type, 1000 replica searches were performed. The results statistics are displayed as box and whiskers plots of percentage of correct prediction (*i*.*e*., correct classification of the variant as evading or non-evading for the considered antibody). The average success in variant classification at the 5^th^ iteration is generally higher than 80%, even ∼100% in some cases. **b**, Illustration of a simple approach to assess the reliability of the prediction (*i*.*e*., the classification as evading or non-evading) for each given sequence. An ensemble of five XGBoost models was used to search the RBD sequence space for evasion/no-evasion of the three antibodies studied: LY-CoV016, LY-CoV555, REGN10987. An evasion score between 0 and 5 was assigned to each sequence as the number of times the variant is predicted to evade the antibody at the fifth iteration. For the three antibodies, most variants are assigned scores of 5 and 0. With very few exceptions, variants with score 5 are found experimentally to evade the antibody, while variants with score 0 do not evade the antibody, again with very few exceptions.

Predictions can never be perfect and must be taken with due caution, in particular when dealing with predictions on antibody evasion. Therefore, it is of interest in this context to propose methods to derive a metric of the reliability of the prediction (*i*.*e*., the classification as evading or non-evading) for each given sequence. Fig. 5b illustrates a simple, practical approach to achieve this. An ensemble of five XGBoost was used to search the RBD sequence space for evasion/no-evasion of the three antibodies studied: LY-CoV016, LY-CoV555, REGN10987. An evasion score between 0 and 5 was assigned to each sequence as the number of times the sequence is predicted to evade the antibody at the fifth iteration. The approach, therefore, involves a total of 5×120=600 sequences in the training set. With very few exceptions (see Fig. 5b), all sequences with score 5 actually evade the antibody and all sequences with score 0 do not evade the antibody. The analysis may also provide an assessment of the overall reliability of the classification for each given antibody, in terms of the number of sequences with intermediate scores (1-4). This number again follows the ranking LY-CoV016 > REGN10987 > LY-CoV555 (Fig. 5b).

## DISCUSSION

We have provided evidence that libraries of, at least, a few hundred thousand variants can be efficiently searched for much enhanced protein properties using state-of-the-art AI tools that are iteratively trained on a much smaller number of variants. To this end, we analysed fitness data on a library of ∼260000 variants of the enzyme dihydrofolate reductase, which have been recently determined using a specialized methodology based on *in vivo* selection.^18^ Our approach identifies the scarce, high-fitness variants of the enzyme from iterative training based on just a few hundred variants. This is a remarkable result because of the prevalence of epistasis in the data,^18^ including higher-order epistasis (*i*.*e*., non-additivity of the effects of more than two mutations), which results in a rugged and challenging fitness landscape to search.

To test the generality of our proposed approach, we applied it to a completely different problem: the emergence of the receptor binding domain (RBD) of the Omicron BA.1 strain of the SARS-CoV-2 virus from that of the parent Wuhan Hu-1 strain. We took advantage of the fact that, recently,^23^ specialized methodologies have been used to determine antibody affinities to essentially all RBD intermediates between the parent and the Omicron strain, *i*.*e*., a library of ∼30000 variants. Despite the fact that pervasive epistasis determines antibody affinity in this case,^23^ we find that, upon iterative training on a total of just a few hundred of variants, the AI tools correctly assign most variants to the evasion or non-evasion classes. This result supports the potential of our approach for the experimental assessment of how viral proteins may evolve in response to antiviral strategies.

The approach we have demonstrated in this work is efficient, first in terms of the computational effort involved, as each search takes only a few seconds on a CPU-based personal computer. It is important to note, furthermore, that we used the default parameters of the AI tools employed (see Methods for details). That is, we did not optimize the AI tools for the specific problems addressed, which further supports the generality of the approach proposed here.

Most importantly, the approach we have developed in this work is also efficient in terms of the number of variants required for iterative training (“learning”). The training sets in the simulations reported here were thus increased by 20 variants at each iteration and, in most cases, the optimization target was already met at the 5^th^ iteration. Also, some analyses were based on ensembles of five, independently trained AI tools, involving a total 100 variants per iteration. We actually selected these numbers (20 and 20×5) on purpose, because they correspond to numbers of variants that can be experimentally studied in the lab, even if specialized methodologies for high-throughput screening are not available. For instance, sets of 100 gene fragments of specified sequences can be ordered from various companies, subsequently cloned and screened as a small library for the desired enzyme property.

Therefore, this work provides a basis for efficient, widely applicable, low-throughput experimental approaches to enzyme optimization.

## METHODS

### Searching the sequence space of the enzyme dihydrofolate reductase

#### Data Preprocessing

Codon sequences and their corresponding fitness values were extracted from the Zenodo’s repository: https://zenodo.org/records/8229020. The dataset fitness_data_wt.csv contains two key columns: **SV**, which included codon sequences, and **m**, representing fitness values. Each codon sequence was transformed into a one-hot encoded numerical representation to facilitate model training.

A one-hot encoding technique was applied to the gene sequences for them to be used by the AI tools selected. The encoding was performed as follows: the gene sequence at the three variable amino acid positions, *i*.*e*., a sequence of 9 consecutive nucleotide positions, was represented as a binary matrix of size 4 × 9. Each column corresponded to one of the four possible nucleotide bases (A, C, G, T) at a specific position. The binary matrix was then flattened into a 1D array of length 36 for compatibility with our computational models.

#### Training and Iterative Enrichment

A Random Forest Regressor, a XGBoost Regressor and a Multi-layer Perceptron Regressor model were employed for predicting fitness values. In each case, the model was trained iteratively over 20 iterations, with 20 initial randomly selected training sequences and 20 additional sequences added at each iteration. The iterative process was conducted as follows: 1) Model Initialization and Training: At each iteration, the Regressor model was trained on the current training dataset, consisting of one-hot encoded nucleotide sequences and their corresponding experimental fitness values; 2) Prediction and Selection: Fitness predictions were made for all sequences in the validation subset. The top 20 sequences with the highest predicted fitness values were selected and added to the training dataset.

#### Replica runs

To explore robustness, searches in sequence space were repeated 1000 times. Each run utilized a different random seed for splitting the dataset into training and validation subsets. After each run, the trained model was saved using the joblib library for further analysis.

#### Implementation details

All computations were conducted using Python. The following libraries were utilized: Pandas^26^ for data manipulation, NumPy^27^ for numerical operations, Scikit-learn^28^ for model design and performance evaluation, Joblib for model serialization and XGBoost package

The XGBoost model was implemented using the XGBRegressor function from the XGBoost library using default parameters except for *n_estimators* (number of trees) that was set to 20.

The Random Forest model was implemented using the RandomForestRegressor function from the scikit-learn library using default parameters except for *n_estimators* (number of trees) that was set to 20.

The Multi-layer Perceptron model was implemented using the MLPRegressor function from the scikit-learn library using default parameters except for hidden_layer_sizes where only one hidden layer was used, and its dimensionality was set to 10. The maximum number of epochs was set at 200 using 0.0001 as tolerance in the optimization process. We used Adam^29^ as solver for the weight’s optimization and ReLU as activation function.^30^ Mean squared error was used as the loss function

For all models, a single run takes a few seconds using a laptop equipped with a 13th Gen Intel® Core™ i7-13700H CPU.

### Searching the sequence space of the receptor binding domain of SARS-CoV-2

#### Data Preprocessing

The dataset used in this study was obtained from the GitHub’s repository: https://github.com/desai-lab/omicron_ab_landscape that accompanies the paper Molulana et al, (2023). In that repository the following files are found: lasso_prediction_REGN10987_aic_opt_order_biochem.txt lasso_prediction_CoV555_aic_opt_order_biochem.txt lasso_prediction_CB6_aic_opt_order_biochem.txt,

We extracted the data from the columns Genotype and Actual Kd in these files. The Genotype column represents the protein sequence encoded in a 15-element vector, where a value of 1 in an element means that the Omicron strain amino acid is present in the corresponding position, while having a 0 means that the amino acid in the RBD of the Wuhan Hu-1 strain is present. The corresponding value in the Actual Kd column is their experimental value for -log(Kd).

To classify genotypes into two classes, we used the following criteria: 1) Genotypes with -log(Kd) values greater than 6.0 were assigned to Class 1 (no evasion). This is equivalent to an antibody dissociation constant smaller than 1 micromolar. 2) Genotypes with -log(Kd) values less than or equal to 6.0 or NaN (Not a Number) were assigned to Class 0 (evasion). This is equivalent to an antibody dissociation constant higher than 1 micromolar.

#### Computational Design

The analysis was conducted across 500 independent runs to explore the statistical robustness of the analysis. Each individual run involved the following steps: 1) Initial Training and Validation Sets: The dataset was randomly split into an initial training set of 20 genotypes and a validation set containing the remaining genotypes. **2)** Iterative Training Process: The iterative learning process was conducted for 20 iterations. During each iteration: a) A classifier model was trained. B) Model Evaluation: The classifier’s accuracy was measured on the validation set. C) Prediction of All Genotypes: The classifier predicted the Class labels for the entire dataset in each iteration.

#### Genotype Selection for Training

To iteratively expand the training set while maintaining diversity we used: 1) Hamming Distance Constraint: a) A Hamming distance threshold was enforced to ensure that new genotypes added to the training set were sufficiently different from those already included. Initially, the threshold was set to 3. Note that, in this context, the Hamming distance^31^ is equivalent to the number of mutational changes. b) If fewer than 20 genotypes met this criterion after 100.000 attempts, the threshold was relaxed to 2, and the search continued. 2) Random Selection: If the above steps failed to identify 20 genotypes, the remaining genotypes were randomly selected from the validation set.

#### Implementation Details

The XGBoost model was implemented using the XGBClassifier function from the XGBoost library using default parameters except for n_estimators (number of trees) that was set to 20.

The Random Forest model was implemented using the RandomForestClassifier function from the scikit-learn library^28^ using default parameters except for *n_estimators* (number of trees) that was set to 20.

The Multi-layer Perceptron model was implemented using the scikit-learn library’s MLPClassifier function using default parameters except for hidden_layer_sizes where only one hidden layer was used and its dimensionality was set to 5. The maximum number of epochs was set at 200 using 0.0001 as tolerance in the optimization process. We used Adam^29^ as solver for the weight’s optimization and ReLU as activation function for the hidden layer. A logistic sigmoid function was used in order to set the binary classification.

All pipelines were written in Python. Key libraries included pandas (McKinney, 2010) for data manipulation, numpy^27^ for numerical operations, and scikit-learn for data splitting and metric computation. Random seeds were set in each run to ensure reproducibility.

For all models, a single run takes a few seconds using a laptop equipped with a 13th Gen Intel® Core™ i7-13700H CPU.

## Code availability

Analyses were performed with custom code deposited with Github: https://github.com/isasio/Virus-Enzyme-Prediction

## Acknowledgments

This research was supported by Grant IHRC22/00004 (to J.M.S.-R.) funded by the “Instituto de Salud Carlos III (ISCIII)” and Next Generation EU, Grant PID2021-124534OB-100 (to J.M. S.-R.) funded by MICIU/AEI/10.13039/501100011033 and Grant PID20210125017OB-I00 (to R.R.-Z.), funded by MCIN/AEI/10.13039/501100011033. The authors would like to thank the DaSCI Institute and the Supercomputing Services of the University of Granada for allowing us to make use of GPU and CPU clusters for the early stages of this work.

## Author contributions

Conceptualization: I.S.-M., R.R.-Z., J.M.S.-R.; Methodology: V.A.R., I.S.-M.; Investigation: I.S.-M.; Funding acquisition: R.R.-Z., J.M.S.-R.; Project administration: J.M.S.-R. Supervision: V.A.R., R.R.-Z., J.M.S.-R.; Writing – original draft: J.M.S.-R.; Writing – review & editing: I.S.-M., V.A.R., R.R.-Z.

## Notes

https://github.com/isasio/Virus-Enzyme-Prediction

### Competing Interest Statement

The authors have declared no competing interest.

